# Improving long-read consensus sequencing accuracy with deep learning

**DOI:** 10.1101/2021.06.28.450238

**Authors:** Avantika Lal, Michael Brown, Rahul Mohan, Joyjit Daw, James Drake, Johnny Israeli

## Abstract

The PacBio HiFi sequencing technology combines less accurate, multi-read passes from the same molecule (subreads) to yield consensus sequencing reads that are both long (averaging 10-25 kb) and highly accurate. However, these reads can retain residual sequencing error, predominantly insertions or deletions at homopolymeric regions. Here, we train deep learning models to polish HiFi reads by recognizing and correcting sequencing errors. We show that our models are effective at reducing these errors by 25-40% in HiFi reads from human as well as *E. coli* genomes.

## Introduction

The PacBio HiFi sequencing method is based upon Single-Molecule, Real-Time (SMRT) sequencing of a circularized DNA molecule in repeated passes, producing multiple subreads [1] with a per-subread accuracy of approximately 90%. Consensus reads are generated using the circular consensus sequencing (CCS) algorithm that includes generation of a graph-based draft consensus using subreads followed by polishing with a Hidden Markov Model. Consensus reads that have a predicted read quality of >Q20 (or less than 1% error) are considered “HiFi reads”.

HiFi sequencing was previously applied to sequence the well-characterized human genome HG002 / NA24385, with an average read length of 13.5 kb and a reported average read accuracy of Q27 (99.8%), approaching that of short-read technologies [2]. This accuracy resulted in improved variant calling performance, approaching that of short reads for small variants while covering a larger proportion of the genome. However, homopolymer errors remain difficult to correct, with 92% of remaining HiFi read errors being indels in homopolymer regions. Since then, HiFi reads have been used in combination with other technologies to produce improved genome assemblies [3–5] as well as improved variant calling performance in a recent precisionFDA challenge [6].

Deep neural networks have recently been used to improve long-read variant calling [7] and to polish assemblies generated from long reads [8]. Here, we train deep learning models to take the pileup of subreads, identify and correct errors in the resulting HiFi reads produced by the CCS algorithm. We show that a neural network architecture with multiple convolutional layers followed by a recurrent layer to integrate long-range sequence information can be trained to reduce CCS errors by 25-40% across multiple HiFi datasets.

## Results

### Deep Learning Improves Consensus Read Accuracy

We obtained previously sequenced human (HG002) sequencing data from [2]. We generated HiFi reads from the subreads using the PacBio CCS algorithm. Each HiFi read was aligned to the human reference genome. For our experiment, we selected reads that aligned within benchmark regions as defined by the Genome In A Bottle (GIAB) consortium [9]. Further, we selected reads that did not overlap with any benchmarked variants within these regions. This was done to eliminate individual-specific genomic variation, so that any remaining differences between the read and the reference sequence could be assumed to be due to sequencing error.

These criteria resulted in a filtered dataset with 27,358 HiFi reads generated from 399,036 subreads; an average of 14.6 subreads per consensus read. For each selected read, the sub-sequence of the human reference genome to which it aligned was extracted and assigned as the label or true sequence.

Reads aligned to chromosome 11 were used for cross-validation. Reads aligning to chromosomes 6, 14 and 20 were used as a test set, while reads aligning to all the remaining autosomes were used for training. There were 22,315 reads in the training set, 1,220 in the validation set, and 3,799 in the test set (Supplementary Table 1).

For each read, a pileup was created by aligning the subreads to the HiFi read. The resulting pileup was then encoded using a summary encoding (Figure 1A).

**Figure 1.**
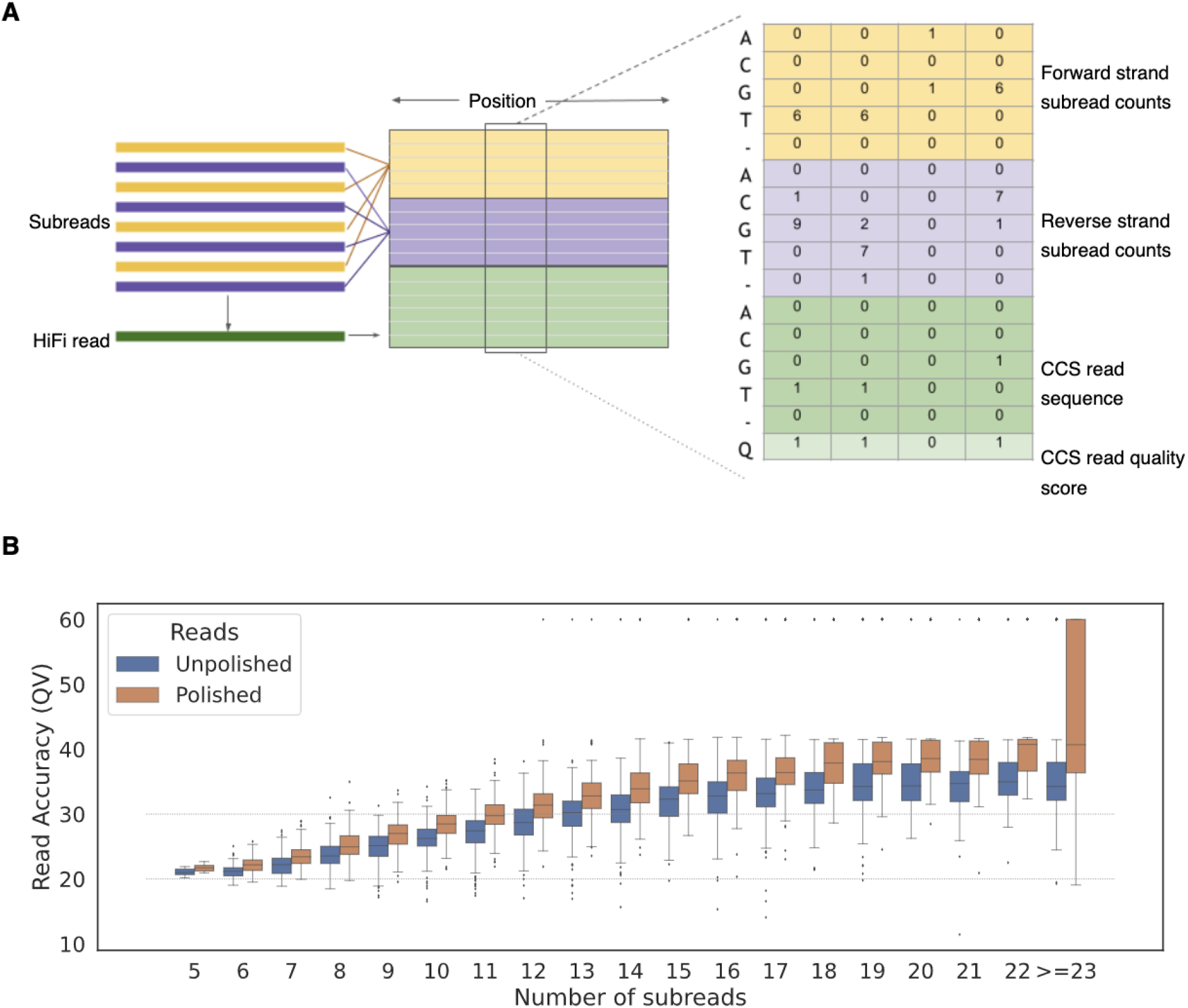
Summary Encoding and Sequel results. **A.** The CCS algorithm is used to generate a HiFi read (green) from forward strand (yellow) and reverse strand (purple) subreads. Both the subreads and the HiFi read are encoded in a matrix of dimensions (16 x no. of positions) where rows 1-5 rows contain the count of bases at each position in the forward strand subreads, rows 6-10 contain the count of bases at each position in the reverse strand subreads, rows 11-15 contain the one-hot encoded HiFi read sequence, and row 16 contains the quality score of each consensus base. **B.** QV read accuracy for unpolished and polished HiFi reads with different numbers of subreads. QV is calculated as −10log_10_(probability of error) and capped at 60. Dotted lines mark QV=20 and QV=30.

For a given pileup, we generated a tensor for each pileup column. We then counted the number of DNA bases (A, C, G, T, deletion) for each pileup column on both the forward and reverse strands. Insertions were handled by encoding new pileup columns. In addition, we encoded the consensus base at each position, and the base quality score for that consensus base. Thus, the encoded pileup is a tensor of shape (number of pileup columns x 16). Of the 16 columns, 5 encode the count of bases on the forward strand, 5 encode the count of bases on the reverse strand, 5 encode the consensus base, and 1 encodes the base quality for the consensus base. Before passing them into the model, the encoded pileup summaries were broken into chunks of length 1000 pileup columns, overlapping by 200 pileup columns. Thus, the model sees examples of shape (16 x 1000).

As this encoding results in loss of information specific to each individual subread, we also attempted to use the complete one-hot encoded pileup of all subreads as an input to our deep learning models. However, our experiments with the complete pileup encoding failed to improve read quality. All results presented below are therefore based on the summary encoding.

We compared several different deep learning model architectures to correct errors in HiFi reads. We initially used Gated Recurrent Unit (GRU) models consisting of two bidirectional GRU layers with a variable number of hidden units; this architecture is similar to that used in previous deep learning models applied to long reads [7,8]. Later, we attempted to reduce the number of model parameters by replacing the first bidirectional GRU layer with convolutional layers.

All models output a softmax activation over five outputs (A, G, C, T, -) at each position of the pileup. The base with the highest softmax value was predicted at each position. A polished HiFi read was produced by combining the predictions for all of the overlapping chunks belonging to the same HiFi read.

While all tested models were close in terms of validation loss, we selected a model with two convolutional layers followed by a bidirectional GRU layer (Supplementary Table 2) for further experiments. When comparing this model to the best bidirectional GRU model, we observed that the model with convolutional layers achieved almost the same validation loss while reducing the number of parameters by 56% and reducing the training time by 68%.

Using this architecture, we found that normalizing the base counts in the reads to sum to 1 slightly reduced validation performance, suggesting that the model benefits from encoding raw counts that indicate the number of subreads at each position. We also observed that the model benefits from access to the HiFi read sequence and quality scores (Supplementary Table 2).

We applied the trained model with the best validation set performance to the 3,799 HiFi reads in the test set (chromosomes 6, 14, and 20). Model inference and writing the corrected reads to a fastq file took 1 hour and 24 minutes.

The corrected reads were aligned to the reference genome and the number of differences between each read sequence and the corresponding reference sequence were counted. We found that the deep learning model improved average HiFi read accuracy from 99.7491% to 99.8461%, reduced the number of errors by 38.7%, and increased the fraction of HiFi reads with QV >= 30 from 0.4485 to 0.5896 (Table 1). The improvement in HiFi read quality is noticeable irrespective of the number of subreads (Figure 1B). Among the 1,080 HiFi reads with 10-15 subreads, the fraction of reads with QV >= 30 increased from 0.451 to 0.751.

**Table 1:** Deep Learning improves HiFi read quality in held-out Sequel HG002 data. DL: Deep Learning. Number of errors was measured as the edit distance between the HiFi read sequence and the reference genome sequence. Read accuracy was measured as (1-(number of errors/alignment length)).

|  | Percent Read Accuracy (mean) | Number of errors per read (mean) | Fraction of reads with QV > 30 |
| --- | --- | --- | --- |
| HiFi | 99.7491 | 32.7823 | 0.4485 |
| HiFi + DL | 99.8461 | 20.1048 | 0.5896 |

### Polishing Sequel II data with deep learning

Since the previous dataset was sequenced using an older Sequel instrument, we next obtained HiFi reads and subreads for an HG002 genome sequenced on a Sequel II system. We processed this dataset in the same way as the Sequel dataset described above, and prepared training, validation and test sets based on the same chromosome splits. The test set consisted of 3,586 HiFi reads from chromosomes 6, 14, and 20.

The baseline quality of this dataset was considerably higher than that of the Sequel dataset (the median QV of HiFi reads in the test set was 32 vs 29 in the Sequel dataset). Nevertheless, we found that the best performing model trained on the Sequel data successfully generalized to this dataset, reducing the number of errors in the test set by 15.9%. However, a model trained using the Sequel II training dataset performed better, reducing errors by 29.6% and increasing average read accuracy from 99.8314 to 99.8821 (Figure 2A, Table 2).

**Figure 2.**
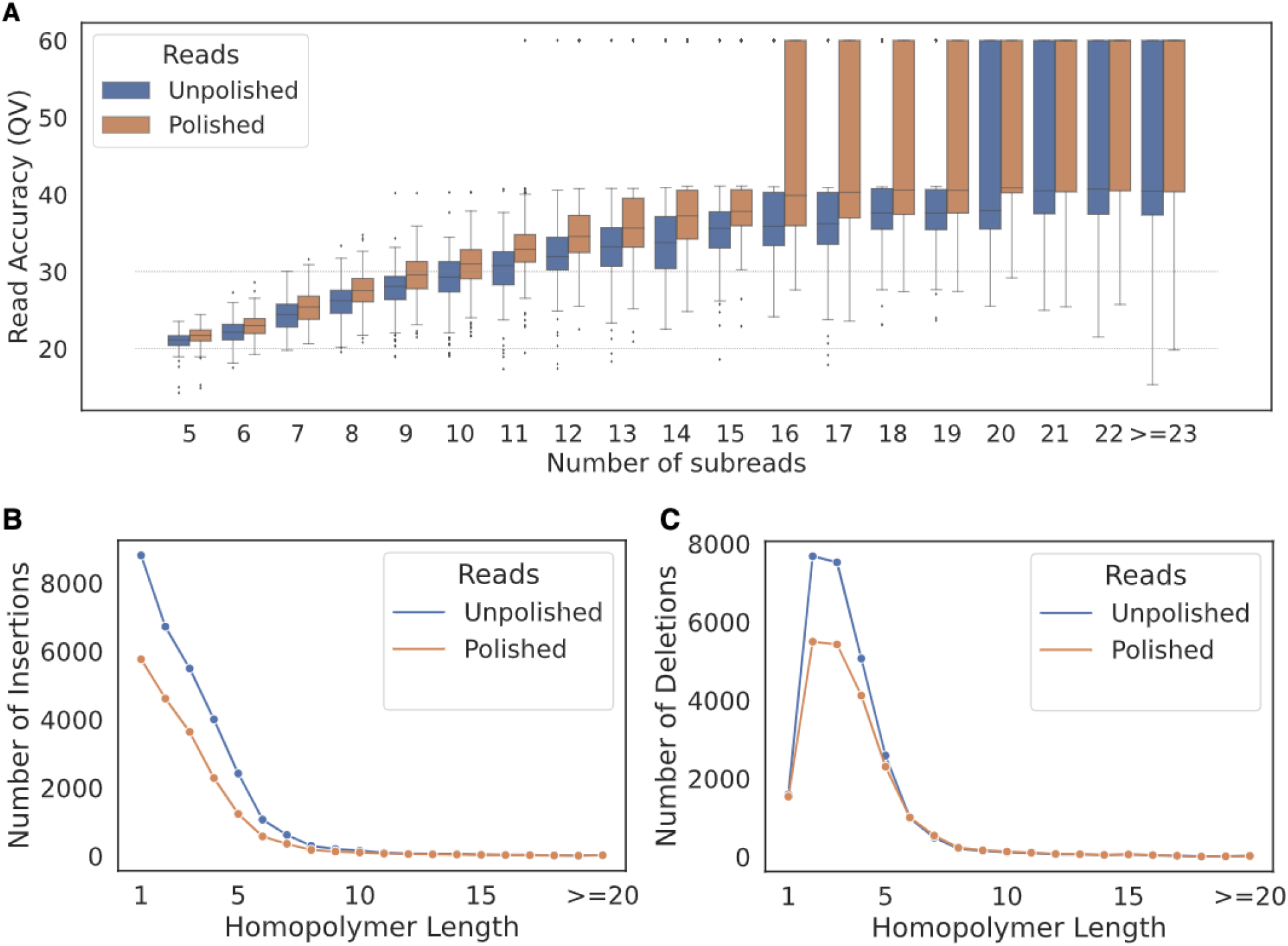
Sequel II HG002 polishing results. **A.** QV Read accuracy for HiFi reads with different numbers of subreads. QV is calculated as −10log_10_(probability of error) and capped at 60. Dotted lines mark QV=20 and QV=30. **B.** Number of insertion errors in HiFi reads before and after polishing, broken down by homopolymer length. **C.** Number of deletion errors in HiFi reads before and after polishing, broken down by homopolymer length.

**Table 2:** Model performance on Sequel II data test sets (HG002 and *E. coli*). DL: Deep Learning. Number of errors was measured as the edit distance between the HiFi read sequence and the true genome sequence. Read accuracy was measured as (1-(number of errors/alignment length)).

|  | Read Accuracy (mean) | Number of errors per read (mean) | Fraction of reads with QV > 30 |
| --- | --- | --- | --- |
| Dataset : HG002 Sequel II |  |  |  |
| HiFi | 99.8314 | 18.5784 | 0.6121 |
| HiFi + DL (trained on Sequel data) | 99.8590 | 15.6319 | 0.6684 |
| HiFi + DL (trained on Sequel II HG002 data) | 99.8821 | 13.0806 | 0.7064 |
| Dataset : *E. coli* Sequel II |  |  |  |
| HiFi | 99.7499 | 35.9272 | 0.3070 |
| HiFi + DL (trained on Sequel II HG002 data) | 99.8157 | 26.5259 | 0.4596 |

We broke down errors in the 3,586 test set reads by type. Overall, deletion errors were reduced by 20% (Figure 2B, Supplementary Table 3) and insertion errors by 36% (Figure 2C, Supplementary Table 3). We further examined abnormal events in the HiFi reads, where an abnormal event was defined as a run of CIGAR alignment errors more than 8 bases long (based on an average read QV of 30, the probability of such an event occurring randomly is (1E-3)^8 = 1E-24). The number of such events was reduced from 20 to 0 in the test set.

### Model generalization from the human genome to *E. coli*

Finally, we tested whether our model can generalize across datasets and species by applying the model trained on HG002 Sequel II data above to an *E. coli* HiFi dataset, which was also sequenced on the Sequel II system. 10,275 HiFi reads from *E. coli* were randomly selected as the test set.

The model trained on HG002 data reduced errors in the *E. coli* reads by 26.2% and increased average read accuracy from 99.7499 to 99.8157 (Fig. 3A, Table 2). However, when we examined the errors by type, we observed an 8% increase in deletion errors in the test set (Fig. 3B, Supplementary Table 4). This was compensated by a 43% decrease in insertions (Fig. 3C, Supplementary Table 4). The number of abnormal events in the test set was reduced from 80 to 10.

**Figure 3.**
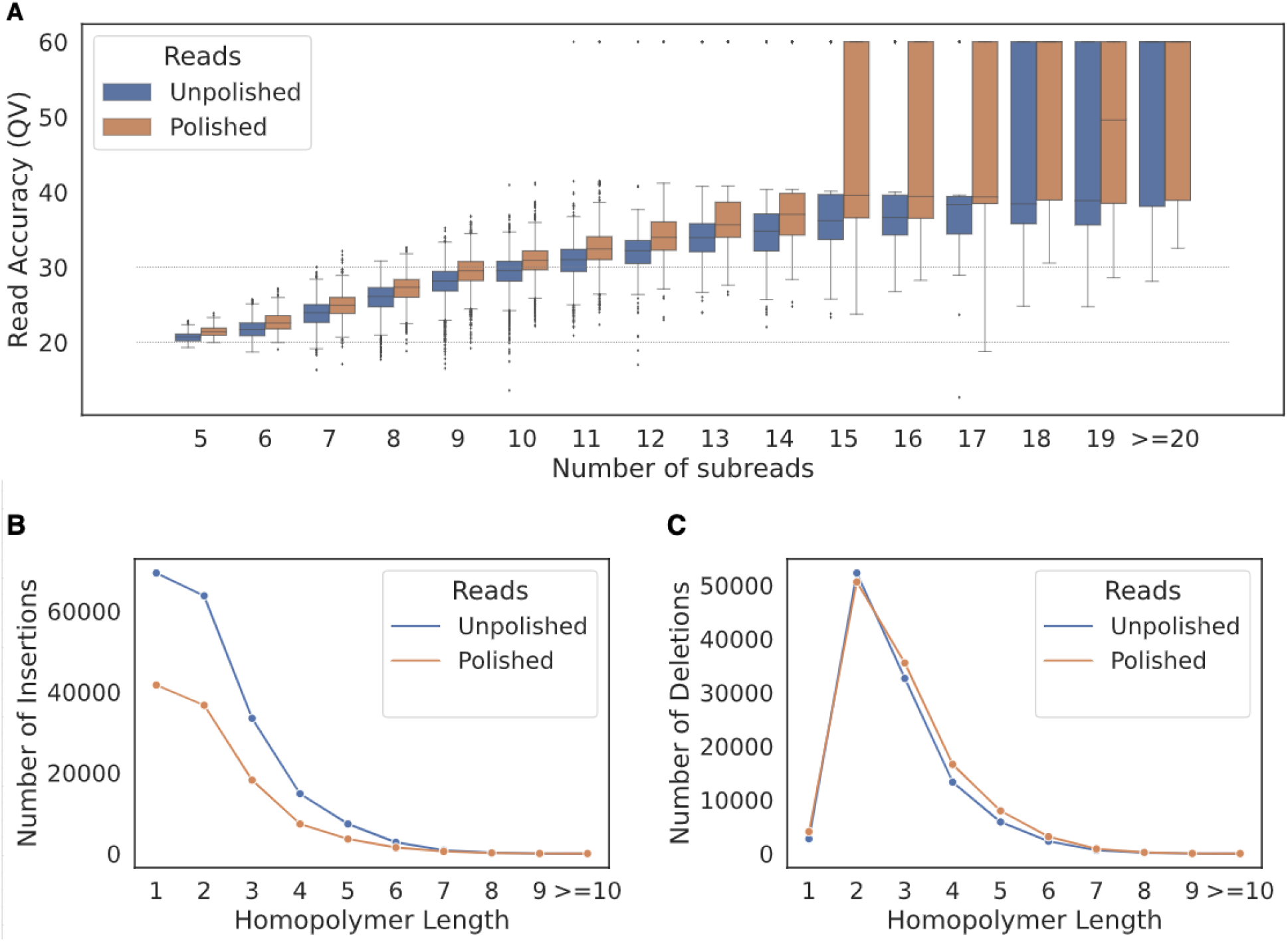
Sequel II *E. coli* polishing results. **A.** QV Read accuracy for HiFi reads with different numbers of subreads. QV is calculated as −10log_10_(probability of error) and capped at 60. Dotted lines mark QV=20 and QV=30. **B.** Number of insertion errors in HiFi reads before and after polishing, broken down by homopolymer length. **C.** Number of deletion errors in HiFi reads before and after polishing, broken down by homopolymer length.

## Discussion

Here we show that deep learning can improve PacBio HiFi read quality, reducing overall error rate by 25-40%. Several different deep learning models were found to be effective at this task. The final model chosen for evaluation consisted of two convolutional layers followed by a bidirectional GRU layer capable of learning long-range features. Our models are able to improve read quality despite not having access to the sequences of individual subreads, or to base-level information from the sequencer such as pulse widths or inter-pulse duration.

We showed that a trained deep learning model corrected insertion and deletion errors in held-out chromosomes of a human (HG002) sequencing dataset. The model was particularly effective at reducing insertion errors that increase homopolymer length and long (>8 bp) stretches of errors. We further showed that a model trained on human data could generalize to a different dataset and species; however, the increase in deletion errors shows that further work is needed to improve generalization. These initial models may be improved by training on more HiFi reads from a larger set of regions in the genome, and by training on data from more diverse sources.

Improving the quality of HiFi reads can help reduce the cost of HiFi sequencing by enabling high-quality consensus reads to be obtained from fewer sequencing passes and consequent shorter run times. In addition, accuracy improvements can improve yield by bringing previously discarded lower quality reads to high quality. Future work may also be directed at enabling more accurate downstream variant calling from these polished long reads, potentially in combination with deep learning models designed for this task [7].

## Methods

### Preparation of human (HG002) sequencing data

For the Sequel HG002 dataset, subreads were downloaded and consensus reads were generated using the CCS algorithm (https://github.com/PacificBiosciences/ccs) version 6.0.0. HiFi reads were aligned to the human reference genome (hs37d5) using pbmm2 version 1.4.0. Alignments falling completely within GIAB (v4.2.1) trusted regions, and not overlapping with any variant, were selected using the bedtools intersect command in bedtools version 2.26.0. For each read, the consensus read sequence, reference sequence (truth) and corresponding subread sequences were extracted. Reads were divided into training, validation and test sets based on the chromosomes to which they aligned.

### Preparation of *E. coli* sequencing data

Subreads and consensus reads were downloaded and aligned to the reference genome using pbmm2 version 1.4.0. Reads overlapping with either of two variant-containing regions were discarded (details are given at https://downloads.pacbcloud.com/public/dataset/Ecoli/egs/) using the bedtools intersect command in bedtools version 2.26.0 [10]. 10,275 reads were randomly chosen from the remaining reads to form the test set.

### Data Encoding

Reads were encoded using the SummaryEncoding class in VariantWorks (https://github.com/clara-parabricks/VariantWorks).

### Models

All deep learning models were implemented using VariantWorks (https://github.com/clara-parabricks/VariantWorks), an open-source, pytorch-based framework to easily train and apply deep learning models to sequencing data. All models were trained for 45 epochs using the Adam optimizer with a learning rate of 0.0001 and a batch size of 256. Experiments were performed using a single Tesla V100 GPU with 16 GB memory.

### Evaluation of results

Polished HiFi reads were evaluated by aligning them to the reference genome using pbmm2 version 1.4.0. Read-level accuracy was extracted from the mc tag in the aligned BAM file. Individual errors at homopolymer regions were extracted and counted using custom scripts.

## Data Availability

HG002 benchmark variants and trusted regions for GRCh37 were downloaded from GIAB version 4.2.1 (https://ftp-trace.ncbi.nlm.nih.gov/ReferenceSamples/giab/release/AshkenazimTrio/HG002_NA24385_son/latest/).

PacBio CCS sequencing data were downloaded from the following sources: Sequel HG002 data: https://downloads.pacbcloud.com/public/publications/2019-HG002-CCS/subreads/

Sequel II HG002 data: https://downloads.pacbcloud.com/public/dataset/HG002_SV_and_SNV_CCS/ Sequel II *E. coli* data: https://downloads.pacbcloud.com/public/dataset/Ecoli/egs/

HiFi reads before and after polishing, along with trained models, are publicly available and can be downloaded from this link: https://nv-pacbio.s3.us-east-2.amazonaws.com/paper_results.zip

## Code Availability

The code to prepare data, train and test deep learning models for CCS read polishing is available at https://github.com/clara-parabricks/VariantWorks/tree/dev-v0.1.0/samples/simple_consensus_caller

## Data availability

HiFi reads before and after polishing are publicly available at https://nv-pacbio.s3.us-east-2.amazonaws.com, along with trained models.

