## Supplementary Tables for "Improving long-read consensus sequencing accuracy with deep learning"

| | Train | Validation | Test |
|---|---|---|---|
| **Chromosomes** | All autosomes except 6, 11, 14, 20 | 11 | 6, 14, 20 |
| **Consensus reads** | 22315 | 1220 | 3799 |
| **Subreads** | 325062 | 18479 | 53222 |
| **Subreads per HiFi read (mean)** | 14.6 | 15.1 | 14.0 |
| **Subreads per HiFi read (median)** | 13 | 13 | 13 |
| **HiFi read length (mean)** | 13042.5 | 13052.2 | 13114.8 |
| **HiFi read length (median)** | 13102.0 | 13081.5 | 13119.0 |
| **HiFi read accuracy (mean)** | 99.7445 | 99.7493 | 99.7491 |
| **HiFi read accuracy (median)** | 99.8767 | 99.8799 | 99.8746 |
| **HiFi read errors (mean)** | 32.4051 | 32.6147 | 32.9039 |
| **HiFi read errors (median)** | 16 | 15 | 16 |

**Supplementary Table 1: Summary of Sequel HG002 training, validation and test sets.** Number of errors was measured as the edit distance between the HiFi read sequence and the true genome sequence. Read accuracy was measured as (1- (number of errors/read length)).

| **Model type** | **Hyperparameters** | **Data Encoding** | **Parameters** | **Validation loss** | **Training time** |
|---|---|---|---|---|---|
| GRU | layers = 1, hidden units = 64 | Default | 32133 | 0.0036354 | 03:49:21 |
| GRU | layers = 2, hidden units = 64 | Default | 106629 | 0.0033968 | 06:09:22 |
| GRU | layers = 1, hidden units = 128 | Default | 113413 | 0.0035039 | 04:23:54 |
| **GRU** | **layers = 2, hidden units = 128** | **Default** | **409861** | **0.0032141** | **10:13:31** |
| **GRU** | **Layers = 2, hidden units = 256** | **Default** | **1606149** | **0.0031371** | **19:46:16** |
| Conv-GRU | Conv layers = 1, kernel width = 3, channels = 128, GRU layers = 1, GRU hidden units = 128 | Default | 205701 | 0.0033022 | 05:34:33 |
| Conv-GRU | Conv layers = 2, kernel width = 3, channels = 128, GRU layers = 1, GRU hidden units = 128 | Default | 254981 | 0.0032350 | 06:01:01 |
| Conv-GRU | Conv layers = 1, kernel width = 3, channels = 256, GRU layers = 1, GRU hidden units = 128 | Default | 310277 | 0.0032968 | 07:28:23 |
| **Conv-GRU** | **Conv layers = 2, kernel width = 3, channels = 256, GRU layers = 1, GRU hidden units = 128** | **Default** | **507141** | **0.0031721** | **08:46:49** |
| Conv-GRU | Conv layers = 2, kernel width = 3, channels = 256, GRU layers = 1, GRU hidden units = 128 | Normalized counts | 507141 | 0.0032584 | 08:49:28 |
| Conv-GRU | Conv layers = 2, kernel width = 3, channels = 256, GRU layers = 1, GRU hidden units = 128 | No HiFi read + quality scores | 502533 | 0.0034818 | 08:41:41 |
| Conv-GRU | Conv layers = 1, kernel width = 3, channels = 128, GRU layers = 1, GRU hidden units = 256 | Default | 601733 | 0.0032411 | 10:11:15 |
| Conv-GRU | Conv layers = 2, kernel width = 3, channels = 128, GRU layers = 1, GRU hidden units = 256 | Default | 651013 | 0.0032442 | 10:24:41 |
| Conv-GRU | Conv layers = 1, kernel width = 3, channels = 256, GRU layers = 1, GRU hidden units = 256 | Default | 804613 | 0.0032170 | 11:27:28 |
| Conv-GRU | Conv layers = 2, kernel width = 3, channels = 256, GRU layers = 1, GRU hidden units = 256 | Default | 1001477 | 0.0031835 | 12:37:38 |

**Supplementary Table 2:** Comparison of model architectures and encodings on the Sequel HG002 validation set.

| Type | Base | Length | Frequency (Unpolished reads) | Frequency (polished reads) | Ratio |
|---|---|---|---|---|---|
| Insertion | A | 1 | 2021 | 1317 | 0.65 |
| Insertion | A | 2 | 1401 | 1153 | 0.82 |
| Insertion | A | 3 | 1443 | 1171 | 0.81 |
| Insertion | A | 4 | 1242 | 883 | 0.71 |
| Insertion | A | 5 | 857 | 526 | 0.61 |
| Insertion | A | 6 | 450 | 285 | 0.63 |
| Insertion | A | 7 | 291 | 175 | 0.60 |
| Insertion | A | 8 | 144 | 94 | 0.65 |
| Insertion | A | 9 | 94 | 61 | 0.65 |
| Insertion | A | 10 | 84 | 54 | 0.64 |
| Insertion | A | 11 | 49 | 37 | 0.76 |
| Insertion | A | 12 | 32 | 27 | 0.84 |
| Insertion | A | 13 | 39 | 27 | 0.69 |
| Insertion | A | 14 | 23 | 14 | 0.61 |
| Insertion | A | 15 | 23 | 23 | 1.00 |
| Insertion | A | 16 | 22 | 15 | 0.68 |
| Insertion | A | 17 | 16 | 13 | 0.81 |
| Insertion | A | 18 | 11 | 8 | 0.73 |
| Insertion | A | 19 | 11 | 6 | 0.55 |
| Insertion | A | 20 | 7 | 5 | 0.71 |
| Insertion | A | 21 | 5 | 4 | 0.80 |
| Insertion | A | 22 | 5 | 5 | 1.00 |
| Insertion | A | 28 | 1 | 1 | 1.00 |
| Insertion | C | 1 | 2319 | 1565 | 0.67 |
| Insertion | C | 2 | 1847 | 1175 | 0.64 |
| Insertion | C | 3 | 1280 | 688 | 0.54 |
| Insertion | C | 4 | 773 | 311 | 0.40 |
| Insertion | C | 5 | 324 | 120 | 0.37 |
| Insertion | C | 6 | 95 | 22 | 0.23 |
| Insertion | C | 7 | 20 | 5 | 0.25 |
| Insertion | C | 8 | 5 | 0 | 0.00 |
| Insertion | C | 9 | 4 | 1 | 0.25 |
| Insertion | C | 10 | 1 | 0 | 0.00 |
| Insertion | C | 11 | 1 | 0 | 0.00 |
| Insertion | G | 1 | 2354 | 1572 | 0.67 |

| | | | | | |
|---|---|---|---|---|---|
| Insertion | G | 2 | 1902 | 1120 | 0.59 |
| Insertion | G | 3 | 1360 | 707 | 0.52 |
| Insertion | G | 4 | 851 | 344 | 0.40 |
| Insertion | G | 5 | 368 | 109 | 0.30 |
| Insertion | G | 6 | 93 | 20 | 0.22 |
| Insertion | G | 7 | 19 | 5 | 0.26 |
| Insertion | G | 8 | 8 | 4 | 0.50 |
| Insertion | G | 9 | 0 | 0 | NA |
| Insertion | G | 10 | 2 | 0 | 0.00 |
| Insertion | T | 1 | 2118 | 1314 | 0.62 |
| Insertion | T | 2 | 1572 | 1166 | 0.74 |
| Insertion | T | 3 | 1414 | 1071 | 0.76 |
| Insertion | T | 4 | 1139 | 746 | 0.65 |
| Insertion | T | 5 | 870 | 480 | 0.55 |
| Insertion | T | 6 | 429 | 247 | 0.58 |
| Insertion | T | 7 | 286 | 175 | 0.61 |
| Insertion | T | 8 | 142 | 81 | 0.57 |
| Insertion | T | 9 | 109 | 69 | 0.63 |
| Insertion | T | 10 | 69 | 48 | 0.70 |
| Insertion | T | 11 | 47 | 35 | 0.74 |
| Insertion | T | 12 | 37 | 29 | 0.78 |
| Insertion | T | 13 | 26 | 21 | 0.81 |
| Insertion | T | 14 | 41 | 30 | 0.73 |
| Insertion | T | 15 | 21 | 12 | 0.57 |
| Insertion | T | 16 | 16 | 13 | 0.81 |
| Insertion | T | 17 | 17 | 12 | 0.71 |
| Insertion | T | 18 | 8 | 7 | 0.88 |
| Insertion | T | 19 | 5 | 4 | 0.80 |
| Insertion | T | 20 | 3 | 2 | 0.67 |
| Insertion | T | 21 | 3 | 3 | 1.00 |
| Insertion | T | 22 | 1 | 1 | 1.00 |
| Insertion | T | 23 | 1 | 1 | 1.00 |
| Insertion | T | 24 | 1 | 1 | 1.00 |
| Deletion | A | 1 | 579 | 559 | 0.97 |
| Deletion | A | 2 | 1724 | 1139 | 0.66 |
| Deletion | A | 3 | 1972 | 1238 | 0.63 |
| Deletion | A | 4 | 1745 | 1225 | 0.70 |

| | | | | | |
|---|---|---|---|---|---|
| Deletion | A | 5 | 1026 | 826 | 0.81 |
| Deletion | A | 6 | 435 | 390 | 0.90 |
| Deletion | A | 7 | 239 | 249 | 1.04 |
| Deletion | A | 8 | 113 | 115 | 1.02 |
| Deletion | A | 9 | 76 | 78 | 1.03 |
| Deletion | A | 10 | 59 | 57 | 0.97 |
| Deletion | A | 11 | 52 | 51 | 0.98 |
| Deletion | A | 12 | 25 | 28 | 1.12 |
| Deletion | A | 13 | 36 | 36 | 1.00 |
| Deletion | A | 14 | 22 | 24 | 1.09 |
| Deletion | A | 15 | 34 | 32 | 0.94 |
| Deletion | A | 16 | 25 | 28 | 1.12 |
| Deletion | A | 17 | 14 | 11 | 0.79 |
| Deletion | A | 18 | 6 | 6 | 1.00 |
| Deletion | A | 19 | 10 | 9 | 0.90 |
| Deletion | A | 20 | 4 | 5 | 1.25 |
| Deletion | A | 21 | 5 | 5 | 1.00 |
| Deletion | A | 22 | 4 | 3 | 0.75 |
| Deletion | A | 28 | 1 | 1 | 1.00 |
| Deletion | C | 1 | 259 | 230 | 0.89 |
| Deletion | C | 2 | 2096 | 1468 | 0.70 |
| Deletion | C | 3 | 1693 | 1311 | 0.77 |
| Deletion | C | 4 | 857 | 809 | 0.94 |
| Deletion | C | 5 | 261 | 292 | 1.12 |
| Deletion | C | 6 | 56 | 79 | 1.41 |
| Deletion | C | 7 | 12 | 17 | 1.42 |
| Deletion | C | 8 | 2 | 3 | 1.50 |
| Deletion | C | 9 | 0 | 3 | NA |
| Deletion | C | 10 | 0 | 2 | NA |
| Deletion | C | 11 | 1 | 1 | 1.00 |
| Deletion | G | 1 | 211 | 211 | 1.00 |
| Deletion | G | 2 | 2038 | 1527 | 0.75 |
| Deletion | G | 3 | 1847 | 1494 | 0.81 |
| Deletion | G | 4 | 840 | 851 | 1.01 |
| Deletion | G | 5 | 297 | 347 | 1.17 |
| Deletion | G | 6 | 57 | 84 | 1.47 |
| Deletion | G | 7 | 6 | 15 | 2.50 |

| | | | | | |
|---|---|---|---|---|---|
| Deletion | G | 8 | 2 | 8 | 4.00 |
| Deletion | G | 9 | 2 | 1 | 0.50 |
| Deletion | G | 10 | 1 | 2 | 2.00 |
| Deletion | T | 1 | 547 | 539 | 0.99 |
| Deletion | T | 2 | 1823 | 1364 | 0.75 |
| Deletion | T | 3 | 2014 | 1385 | 0.69 |
| Deletion | T | 4 | 1625 | 1236 | 0.76 |
| Deletion | T | 5 | 1003 | 837 | 0.83 |
| Deletion | T | 6 | 438 | 452 | 1.03 |
| Deletion | T | 7 | 235 | 262 | 1.11 |
| Deletion | T | 8 | 93 | 107 | 1.15 |
| Deletion | T | 9 | 70 | 83 | 1.19 |
| Deletion | T | 10 | 53 | 72 | 1.36 |
| Deletion | T | 11 | 43 | 52 | 1.21 |
| Deletion | T | 12 | 35 | 47 | 1.34 |
| Deletion | T | 13 | 25 | 31 | 1.24 |
| Deletion | T | 14 | 22 | 25 | 1.14 |
| Deletion | T | 15 | 26 | 28 | 1.08 |
| Deletion | T | 16 | 17 | 16 | 0.94 |
| Deletion | T | 17 | 13 | 17 | 1.31 |
| Deletion | T | 18 | 5 | 2 | 0.40 |
| Deletion | T | 19 | 5 | 4 | 0.80 |
| Deletion | T | 20 | 1 | 1 | 1.00 |
| Deletion | T | 21 | 3 | 4 | 1.33 |
| Deletion | T | 22 | 1 | 1 | 1.00 |
| Deletion | T | 23 | 1 | 2 | 2.00 |
| Deletion | T | 24 | 1 | 2 | 2.00 |

**Supplementary Table 3:** Breakdown of insertion and deletion errors in the Sequel II HG002 test set.

| Type | Base | Length | Frequency (Unpolished reads) | Frequency (polished reads) | Ratio |
|---|---|---|---|---|---|
| Insertion | A | 1 | 13693 | 7211 | 0.53 |
| Insertion | A | 2 | 10206 | 6247 | 0.61 |
| Insertion | A | 3 | 7309 | 4679 | 0.64 |
| Insertion | A | 4 | 4482 | 2564 | 0.57 |
| Insertion | A | 5 | 2926 | 1645 | 0.56 |
| Insertion | A | 6 | 1297 | 794 | 0.61 |
| Insertion | A | 7 | 386 | 263 | 0.68 |
| Insertion | A | 8 | 92 | 69 | 0.75 |
| Insertion | A | 9 | 12 | 9 | 0.75 |
| Insertion | C | 1 | 20253 | 13506 | 0.67 |
| Insertion | C | 2 | 21104 | 12703 | 0.60 |
| Insertion | C | 3 | 9317 | 4770 | 0.51 |
| Insertion | C | 4 | 2896 | 1332 | 0.46 |
| Insertion | C | 5 | 744 | 307 | 0.41 |
| Insertion | C | 6 | 135 | 56 | 0.41 |
| Insertion | C | 7 | 25 | 9 | 0.36 |
| Insertion | C | 8 | 2 | 0 | 0.00 |
| Insertion | C | 10 | 1 | 0 | 0.00 |
| Insertion | G | 1 | 20662 | 13162 | 0.64 |
| Insertion | G | 2 | 21896 | 11668 | 0.53 |
| Insertion | G | 3 | 9457 | 4548 | 0.48 |
| Insertion | G | 4 | 3036 | 1301 | 0.43 |
| Insertion | G | 5 | 747 | 273 | 0.37 |
| Insertion | G | 6 | 123 | 53 | 0.43 |
| Insertion | G | 7 | 27 | 8 | 0.30 |
| Insertion | G | 8 | 8 | 6 | 0.75 |
| Insertion | G | 9 | 1 | 1 | 1.00 |
| Insertion | T | 1 | 14832 | 7828 | 0.53 |
| Insertion | T | 2 | 10588 | 6083 | 0.57 |
| Insertion | T | 3 | 7391 | 4182 | 0.57 |
| Insertion | T | 4 | 4393 | 2120 | 0.48 |
| Insertion | T | 5 | 2915 | 1357 | 0.47 |
| Insertion | T | 6 | 1225 | 591 | 0.48 |
| Insertion | T | 7 | 386 | 241 | 0.62 |
| Insertion | T | 8 | 105 | 74 | 0.70 |

| | | | | | |
|---|---|---|---|---|---|
| Insertion | T | 9 | 7 | 5 | 0.71 |
| Deletion | A | 1 | 812 | 1337 | 1.65 |
| Deletion | A | 2 | 6881 | 7574 | 1.10 |
| Deletion | A | 3 | 6729 | 7181 | 1.07 |
| Deletion | A | 4 | 3980 | 4623 | 1.16 |
| Deletion | A | 5 | 2379 | 3073 | 1.29 |
| Deletion | A | 6 | 1064 | 1373 | 1.29 |
| Deletion | A | 7 | 286 | 395 | 1.38 |
| Deletion | A | 8 | 70 | 92 | 1.31 |
| Deletion | A | 9 | 8 | 11 | 1.38 |
| Deletion | C | 1 | 598 | 760 | 1.27 |
| Deletion | C | 2 | 19389 | 17103 | 0.88 |
| Deletion | C | 3 | 9688 | 9989 | 1.03 |
| Deletion | C | 4 | 2607 | 3276 | 1.26 |
| Deletion | C | 5 | 577 | 796 | 1.38 |
| Deletion | C | 6 | 102 | 146 | 1.43 |
| Deletion | C | 7 | 18 | 26 | 1.44 |
| Deletion | C | 8 | 1 | 3 | 3.00 |
| Deletion | C | 10 | 1 | 2 | 2.00 |
| Deletion | G | 1 | 553 | 732 | 1.32 |
| Deletion | G | 2 | 19117 | 17946 | 0.94 |
| Deletion | G | 3 | 9590 | 10686 | 1.11 |
| Deletion | G | 4 | 2659 | 3515 | 1.32 |
| Deletion | G | 5 | 636 | 871 | 1.37 |
| Deletion | G | 6 | 105 | 152 | 1.45 |
| Deletion | G | 7 | 24 | 31 | 1.29 |
| Deletion | G | 8 | 5 | 6 | 1.20 |
| Deletion | G | 9 | 1 | 1 | 1.00 |
| Deletion | T | 1 | 783 | 1251 | 1.60 |
| Deletion | T | 2 | 6894 | 7989 | 1.16 |
| Deletion | T | 3 | 6633 | 7662 | 1.16 |
| Deletion | T | 4 | 4067 | 5203 | 1.28 |
| Deletion | T | 5 | 2290 | 3220 | 1.41 |
| Deletion | T | 6 | 1019 | 1470 | 1.44 |
| Deletion | T | 7 | 287 | 423 | 1.47 |
| Deletion | T | 8 | 81 | 112 | 1.38 |
| Deletion | T | 9 | 6 | 7 | 1.17 |

**Supplementary Table 4:** Breakdown of insertion and deletion errors in the Sequel II *E. coli* test set.
